# Respiratory microbiota as a health biomarker in blue, fin and humpback whales: Pilot study in the Gulf of St-Lawrence (Québec, Canada)

**DOI:** 10.64898/2026.03.16.711931

**Authors:** Anik Boileau, Jonathan Blais, Catharina Vendl, Roxane Plante, Marion Desmarchelier, Marcio Costa, André Marette, Kathleen Hunt, Jamie Ahloy-Dallaire

**Affiliations:** Département des Sciences animales, Université Laval, Québec, QC G1V 0A6, Canada; Centre d’Éducation et de Recherche de Sept-Îles, Sept-Îles, QC G4R 5B7, Canada; School of Agricultural, Environmental and Veterinary Sciences, Charles Sturt University, Boorooma NSW 2650, Australia; Faculté de Médecine Vétérinaire, Université de Montréal, Québec, QC J2S 2M2, Canada; Faculté de Médecine, Université Laval, Québec, QC G1V 0A6, Canada; Marine Mammal Institute, Oregon State University, Corvallis, OR 97331

**Keywords:** Rorqual Whales, Pathobionts, Alpha Diversity, Health, Welfare Indicators

## Abstract

Amplicon-based profiling of airway microbiota is increasingly used to assess respiratory health in mammals, yet baseline data for free-ranging baleen whales remain scarce. We characterised the exhaled-breath (“blow”) microbiota of rorqual whales in the Gulf of St. Lawrence (Canada) and examined associations with individual health indicators. Blow samples were collected opportunistically from six whales (two blue, two fin and two humpback), with seawater and air controls. The V4 region of the 16S rRNA gene was sequenced on an Illumina MiSeq platform and processed in R (v4.5) using the DADA2 pipeline for quality filtering, denoising and amplicon sequence variant (ASV) inference. Alpha diversity varied among individuals (Shannon = 2.72 – 4.33) and beta-diversity analyses revealed a significant effect of environment (whale blow vs. seawater) on community composition (PERMANOVA: *R*^2^ = 0.140, *F* = 1.31, *p* = 0.030). The relative abundance of pathobionts (22.8–48.8%) was negatively correlated with alpha diversity (Spearman ρ = −0.88 to −0.94, p < 0.05), while higher diversity correlated positively with good skin condition (ρ = 0.84, p = 0.03). These findings provide the first baseline description of rorqual respiratory microbiota in the Gulf of St. Lawrence and support blow microbiome metrics as non-invasive health biomarkers.

## 2. Introduction

In recent years, studies on the respiratory health of human and non-human animals have highlighted the important role of the airway microbiota composition in the overall homeostasis or dysbiosis of their host’s respiratory system [1–3]. Both the upper and lower airway tract microbiota have been linked to respiratory health and disease in production animals [4–9], companion animals [10–12], laboratory animals [13–15], and humans [16–18]. Important similarities and differences in commensal microbiota have been observed between species and their respective physiology [1,19]. For instance, the upper respiratory tracts (URT) of healthy bovine, porcine, feline and canine species are similar both anatomically and in their commensal microbiota, which are dominated, at the phylum level, by Proteobacteria, Tenericutes, Firmicutes and Bacteroidetes [11,20–26]. Moreover, endogenous respiratory variables, such as pH, osmolarity, and temperature, and extrinsic factors, such as the maternal vaginal microbiome, housing conditions, or pathogen exposure, have also been associated with the respiratory microbiota composition in domestic mammals [27]. Studies have shown that each species has a unique microbial community and that greater alpha diversity of the airway microbiota acts as a “gatekeeper” of its host’s respiratory health [1,16,19]. Conversely, it has been reported that airway microbiota that contain a wide range of pathobionts such as y-Proteobacteria are associated with dysbiosis of the respiratory tract and infectious diseases [28–30].

In cetaceans, extensive differences from terrestrial mammals can be observed both anatomically and environmentally. In odontocetes (toothed whales, dolphins and porpoises) the respiratory and digestive tracts are fully separated. In contrast, baleen whales such as rorquals possess a partially separated respiratory system, characterised by two blowholes connected to the larynx, nasopharynx, laryngeal air sac, trachea and lungs [31]. During feeding in baleen whales, a musculo-fatty structure called “the oral plug” seals the oropharyngeal tract to prevent water from entering the respiratory tract, although it is expected that small amounts of water might still enter the upper portion of the nasal cavity/ blowhole margins during ventilation cycles at the surface [32–34]. Furthermore, cetacean lungs have few mucus-producing cells, no coughing reflexes, and little lymphoid tissue, critical in housing immune cells to fight pathogens [35,36]. These specificities, associated with the marine environment, underline how cetaceans are subjected to not only microbial airborne communities but also to seawater ones concomitantly [37,38].

Studies on the exhaled breath microbiota of cetaceans have been conducted on captive and free-ranging odontocetes [39–45] but also in baleen whales like the gray (*Eschrichtius robustus*) [46], humpback (*Megaptera novaeangliae*) [34,46–49], blue (*Balaenoptera musculus*) [47,50,51], and fin (*Balaenoptera physalus*) whales [52]. Humpback whales appear to exhibit a relatively consistent core respiratory microbiome across years and geographic regions (e.g., Pacific and Atlantic populations), with commonly reported genera including *Bacillus, Burkholderia, Propionibacterium, Anoxybacillus*, and *Geobacillus*. Dominant phyla generally include Actinobacteria, Bacteroidetes, Firmicutes, and several classes of Proteobacteria [34,39,46–49]. Similar taxonomic patterns have also been reported in blue whales from the Gulf of California [47,50,51], whereas in fin whales sampled in the same region, *Haemophilus* spp. was the only bacterial genus identified [52]. Several studies have also reported bacterial taxa previously associated with respiratory pathogens or airway dysbiosis. For example, Vendl et al. (2020) showed that fasting during the annual migration of East Australian humpback whales to breeding grounds was associated with reduced diversity and richness of their exhaled breath microbiota [47]. The study also identified ten core genera previously described as pathogenic in cetaceans [53,54], although their occurrence was not related to the whales’ fasting stage [47]. Similarly, Apprill et al. (2017) detected several genera previously considered potentially pathogenic in marine mammals, although none were linked to active respiratory disease in the individuals examined. Notably, *Corynebacterium* was part of the core respiratory microbiota of all whales in that study [34,53], despite having previously been isolated only from the spleen and brains of clinically ill dolphins with systemic bacterial infections prior to death [53]. Thus, some descriptions of respiratory microbiota for humpback, blue and fin whales have been performed, though as of yet with relatively small sample sizes and from only a few locations. However, no studies have yet been performed in the Gulf of St. Lawrence (Québec, Canada), a well-documented anthropized environment. Further, no studies have yet explored possible relationships between respiratory microbiome and other indices of whale health and welfare, an important data gap given the potential of cetacean respiratory microbiome analysis as a non-invasive metric of general health.

Each year, several species of rorqual whales, including humpback, blue, and fin whales, enter the Gulf of St-Lawrence, (hereafter GSL) in early spring (April-May) to feed for several months. During this period, they exploit a variety of prey resources, including pelagic fishes such as Atlantic herring (*Clupea harengus*) and capelin (*Mallotus villosus*), as well as krill (*Euphausiacea*). These small, shrimp-like crustaceans constitute the exclusive food source of blue whales, which are specialist feeders compared to humpback and fin whales [55– 60]. However, due to growing anthropogenic activities and climate change, studies spanning three decades have highlighted recent shifts in the distribution, abundance, and feeding habits of these species [61–65].

Moreover, a recent study showed that fishing activities and altered seawater parameters can impact the physical health and welfare of the rorqual whales found in this feeding ground [66]. To date, no studies have characterised the respiratory microbiota of the rorqual whales found in the GSL. Considering that the Canadian Species at Risk Act has listed blue and fin whale populations found in the GSL as endangered and of special concern, respectively [67,68], it is crucial to monitor health parameters in these animals. The current study aimed to explore the core respiratory microbiota of humpback, blue, and fin whales in the Gulf of St. Lawrence, a region increasingly impacted by human activities, to provide a baseline assessment for long-term health and welfare assessment in these species. We hypothesised that intra-individual airway microbiota signatures would differ between each whale and the surrounding seawater microbial community. We further aimed to explore the relationship between alpha diversity metrics, the relative abundance of microbial pathogens (confirmed and opportunistic), and other metrics of individual whale health (e.g. body condition, skin condition, etc.) to validate respiratory microbiome biomarkers as a health and/or potential welfare measure that could be included in further health and welfare assessment protocols. Finally, our study takes advantage of a unique sequence of three successive samples of the same individual to examine intra-individual consistency in microbial diversity metrics which, to our knowledge, has never been done before.

## 3. Materials and Methods

Aboard a small research platform (19-foot Brig (2020) with a 90 hpw outboard motor), individual whales were opportunistically located and approached between June and October (2020-2025) in the Sept-Îles area of the Gulf of St-Lawrence (Fig.1). Photographs and video recordings (for whale ID and welfare/physical health assessments [66]), and respiratory vapours for microbiota gene sequencing were collected during approach sequences lasting under 120 minutes for each individual whale. Once an animal came up to the surface to engage in a ventilation sequence, we approached slowly and carefully from either side and swung a 16-foot pole, carrying a petri dish apparatus, back and forth in the air to capture the exhaled breath droplets. (Fig.2). When the whale began a deep dive, the boat stopped, and the petri dishes (*n* = 1-3) were swabbed with a flocked nylon fibre bud (E-Swabs, Copan) and placed in modified Amies medium vial (1 ml) for transport (on ice, for approximately 3 hours), and storage at -80°C. Sea water samples (*n* = 2), from the same area (same day as blow samples), were collected by dipping a petri dish briefly on the sea surface layer, and one air sample was collected out at sea with a petri dish mounted on the pole up in the air when the whales had left the area. Air and seawater samples were transported and stored in the same manner as blow samples until laboratory analysis.

**Figure 1.**
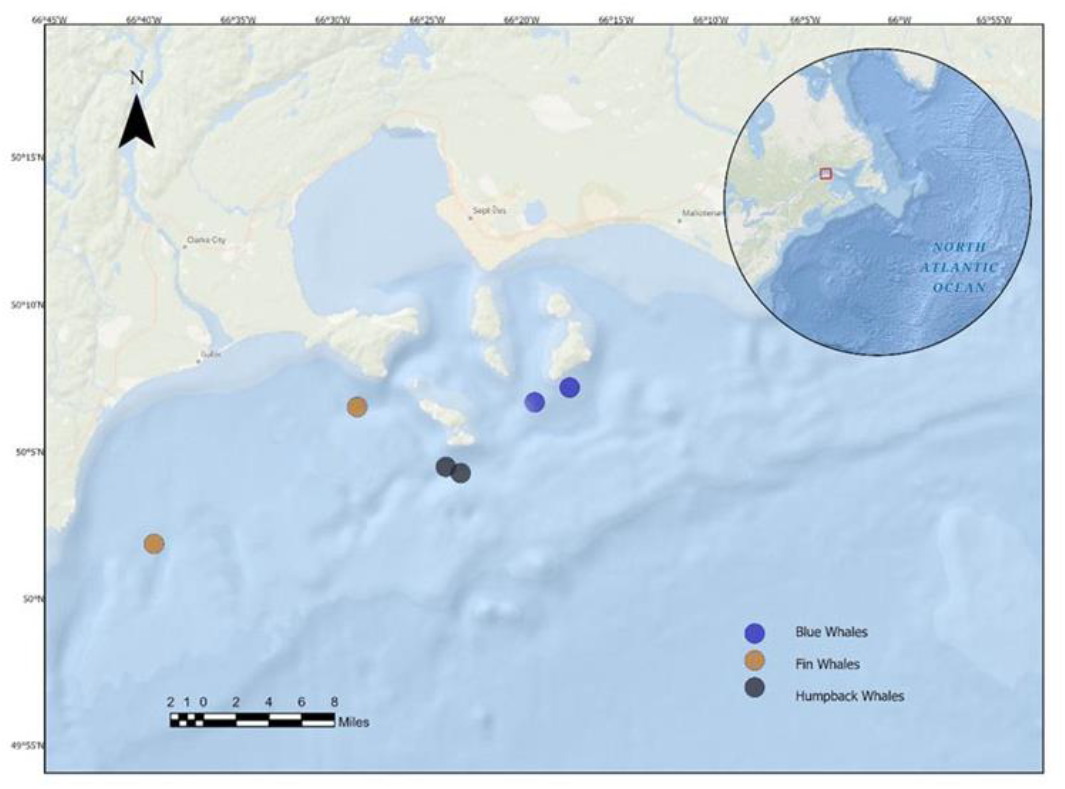
Research area in Sept-Îles, Gulf of St-Lawrence (Québec, Canada), and geographical position of the whales that were sampled.

**Figure 2.**
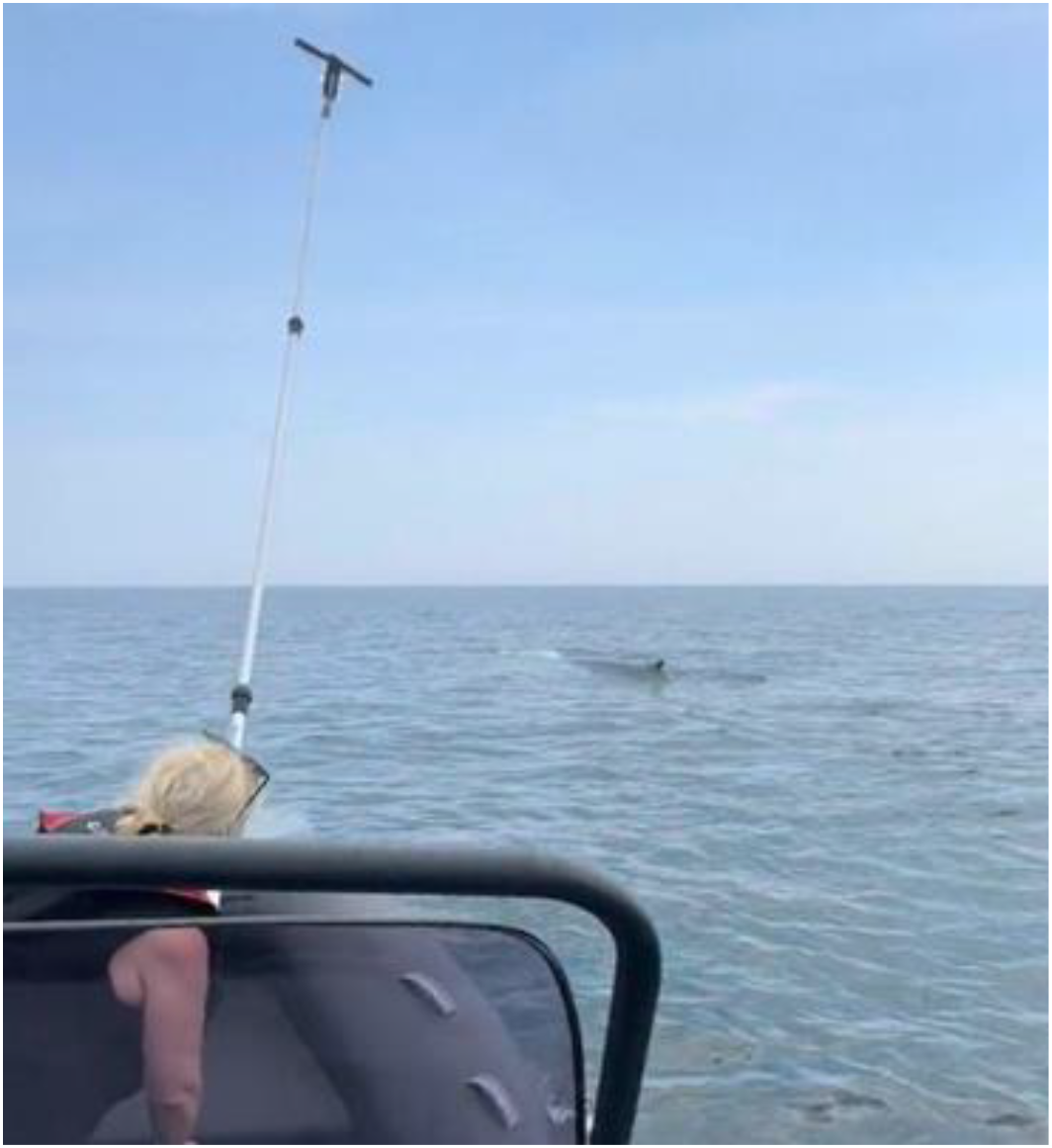
The 16-foot pole with petri dishes attached at the tip, being used to collect respiratory vapours from a blue whale. (Credit André Marette)

The present study (2020-2025) is part of a larger project on the monitoring of health and welfare of rorqual whales found in the GSL (since 2017). The current research is in compliance with the Canadian Council on Animal Care [69] and received ethical approval from the Animal Care Committee of the University Laval, and the Division of Fisheries and Oceans Canada.

### 3.1 DNA extraction and 16S rRNA gene sequencing

Genomic DNA was isolated from respiratory vapour samples using the glass-bead-based DNeasy PowerSoil® Kit (QIAGEN, Toronto, ON, Canada), according to the manufacturer’s instructions. DNA concentration and purity were assessed spectrophotometrically using a NanoDrop™ 1000 (Thermo Fisher Scientific, Waltham, MA, USA), and samples were stored at −20 °C until further analysis. The V4 region of the bacterial 16S rRNA gene was amplified by PCR using primers 515F (5′-GTGCCAGCMGCCGCGGTAA-3′) and 806R (5′-GGACTACHVGGGTWTCTAAT-3′) [42]. Amplicon libraries were then subjected to paired-end sequencing (2 × 250 bp) on an Illumina MiSeq platform (Illumina, San Diego, CA, USA) at the McGill Génome Québec Innovation Centre (Montréal, QC, Canada).

### 3.2 Bioinformatic analysis

FASTQ files generated by Illumina MiSeq sequencing were processed in R (v4.3.3) using the DADA2 software package [70]. Primers were first removed, and following quality assessment, reads were truncated at 225 bp (forward) and 220 bp (reverse) to maximise sequence retention. Reads were then trimmed, dereplicated, denoised and merged to infer high-resolution amplicon sequence variants (ASVs), which distinguish sequences differing by as little as one nucleotide [71]. Chimeric sequences were subsequently identified and removed. Taxonomic assignment for each ASV was performed using a Naive Bayes classifier trained on the SILVA v132 database [72], a curated reference set of aligned 16S rRNA gene sequences widely used for microbial identification. Finally, potential contaminants associated with the blank control were identified and removed using the *decontam* package.

### 3.3 Health measures analysis

Based on the physical indicator welfare assessment protocol developed by Boileau et al. 2024 [66], photographs of the animals were scored with the 0-100 scales for: body condition, skin health condition, injury condition and ectoparasite loads, where the lowest scores reflect the poorest health/welfare states.

### 3.4 Statistical analysis

All statistical analyses were performed in R (v4.3.3). Alpha-diversity metrics (observed richness in ASVs, Shannon–Wiener index, inverse Simpson index, and Faith’s phylogenetic diversity (PD)) were calculated for each sample using the ASV count matrix and corresponding taxonomy table generated by DADA2.

Differences in alpha diversity among samples were assessed using non-parametric Kruskal-Wallis tests, followed by pairwise Wilcoxon rank-sum tests for each metric.

Beta-diversity analyses were conducted to compare the microbial community structure of whale respiratory samples (blow) with air and seawater samples. Community dissimilarities were quantified using Bray-Curtis, Jaccard, and UniFrac (weighted and unweighted) distances. Bray-Curtis dissimilarities were calculated to capture differences in community structure based on taxonomic abundances, whereas Jaccard distances were computed from presence-absence matrices to compare ASV occurrence independent of abundance. For phylogeny-aware comparisons, ASV sequences were aligned using the *DECIPHER* package and a maximum-likelihood phylogenetic tree was inferred using the *phangorn* package; weighted and unweighted UniFrac distances were then computed in *phyloseq*, incorporating phylogenetic branch lengths.

Differences in community composition between whale blow, air and seawater samples were tested using PERMANOVA (adonis2 function in the *vegan* package) with 999 permutations and Group (Whale vs Seawater) as the explanatory variable. Prior to interpreting PERMANOVA results, homogeneity of multivariate dispersion was assessed using betadisper (*vegan*). Ordination analyses were performed using Principal Coordinates Analysis (PCoA) for each distance metric to visualise clustering patterns. Finally, Spearman’s rank correlation tests were used to assess relationships between alpha-diversity metrics (Shannon-Weiner index, inverse Simpson index, and Faith’s PD), health parameters (body condition, skin health, injury condition and ectoparasite load scores) and the ratios of all pathobionts to commensal microbiota in each whale. For all tests, statistical significance was set at α = 0.05.

## 4. Results

Amplicon sequencing of the V4 region generated 286,168 raw 16S rRNA gene reads across 12 samples: two technical controls (sea air and blank petri dish), two seawater samples, and eight whale respiratory blow samples: two from humpbacks (ID Mn050 and Mn051), four from fin whales (three from fin whale ID Bp053 who ventilated 12 times during this one sequence: Bp053-01 = at the beginning of the respiratory sequence after 12 minutes of diving; Bp053-02 = at the middle of the same respiratory sequence, and Bp053-03 = at the end of the respiratory sequence, just before diving for ≥ 10 minutes; and one from fin whale Bp020), and finally two from blue whales (Bm001 and Bm002). After quality filtering, denoising, paired-end merging and chimaera removal, 225,745 reads (78.9%) were retained for downstream analyses. We removed 16 ASVs associated with the blank control, since they were considered contaminants. The resulting dataset contained a total of 1034 ASVs, across 26 phyla, 44 classes, 105 orders, 160 families, and 245 genera. The whale’s life status and physical welfare assessment results showed that the two humpback whales (Mn050 and Mn051) were juveniles (both ≤ 4 years old), compared to Bp053 who was considered an adult (known since 2016), Bp020 known as an adult since 1987, therefore considered an elder, Bm001 known since 1994, also an elder, and blue whale Bm002 known since 2007, so considered at least an adult. Bp053 was considered to be unhealthy; he was thin and had Tattoo-skin-disease-like lesions on his rostrum, his right-side flank and peduncle. The two blue whales were extremely thin, and the two humpbacks were thin and had multiple skin scratches on their entire bodies, likely inflicted by conspecifics.

### 4.1 Alpha diversity

Observed richness (ASVs), diversity evenness (Shannon-Weiner Index), diversity dominance (Inverse-Simpson), and Faith’s phylogenetic diversity (PD) varied among sample types (Table 1). Seawater samples exhibited the highest richness values (155–270 ASVs; SD = 81.31), consistent with the extensive microbial heterogeneity typically found in marine water. These samples also showed relatively high diversity evenness (Shannon: 3.05–4.47, SD = 1,0), dominance (inverse Simpson diversity: 5.83–37.99, SD = 22.7), and Faith’s PD (22.48-17.06; SD = 3.83), indicating both taxonomic breadth and evenness.

**Table 1.**
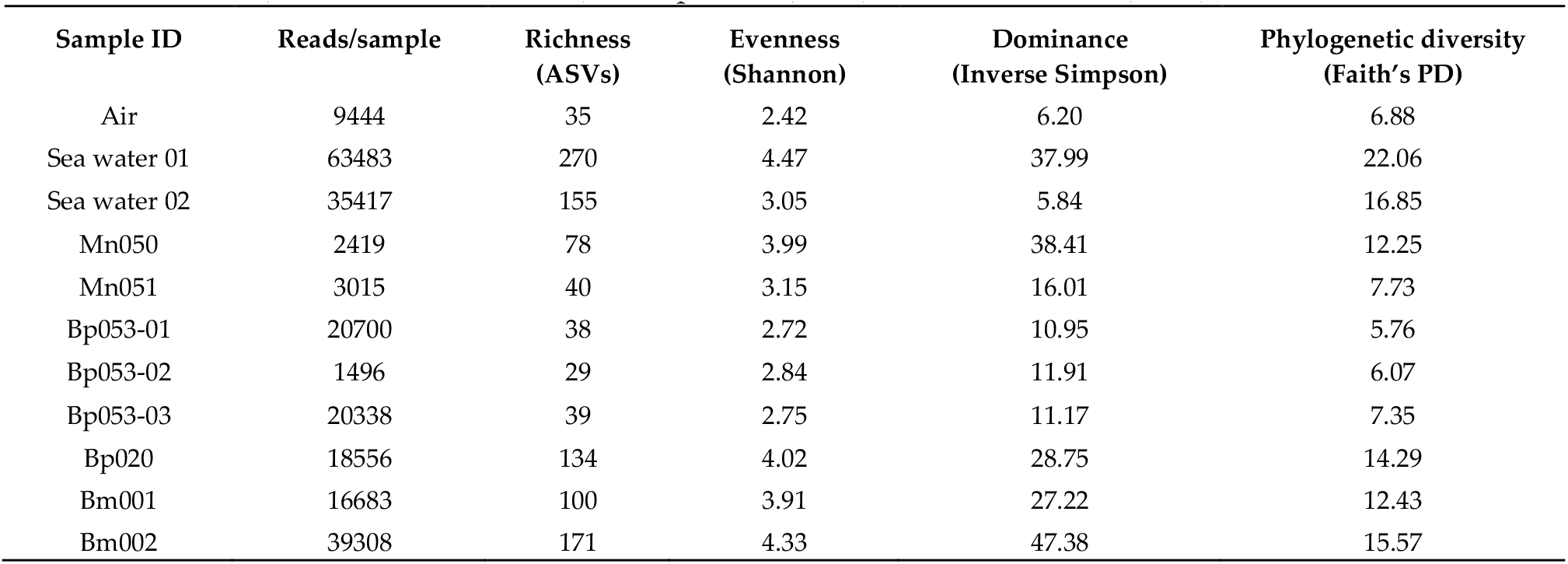
Alpha diversity values and reads per sample for control sample: air (*n* = 1), seawater samples (*n* = 2), and whales: fins (*n* = 4 for 2 individuals), humpbacks (*n* = 2) and blue whales (*n* = 2).

In comparison, whale blow samples showed intermediate and variable alpha diversity, with richness ranging from 28 to 171 ASVs (SD = 100.11), evenness (Shannon-Weiner) from 2.72 to 4.33 (SD = 1.14), dominance (inverse Simpson) ranging from 10.95 to 47.38 (SD = 25.76), and phylogenetic diversity (Faith’s PD) ranging from 5.7 to 15.57 (SD = 6.98). The air control sample displayed the lowest richness and diversity (35 ASVs; Shannon-Weiner = 2.42; inverse-Simpson = 6.20, Faith’s PD= 6.65), demonstrating minimal background amplification.

Alpha diversity did not differ significantly between air control and whale samples for ASV richness, Shannon-Weiner evenness, inverse Simpson dominance, or Faith’s phylogenetic diversity (all *p* > 0.12). Comparisons between seawater and whale samples revealed no significant differences in taxonomic alpha diversity; however, Faith’s phylogenetic diversity was significantly lower in whale samples compared to seawater (Kruskal-Wallis, *χ*^2^ = 4.36, *p* = 0.037; pairwise Wilcoxon, *p* = 0.044). No significant differences were detected in any alpha diversity metric between whale species (fin, humpback, and blue; all *p* > 0.20) (Table 2).

**Table 2.** Alpha diversity comparison between all samples according to Kruskal-Wallis and post-hoc Pairwise Wilcoxon with Benjamini–Hochberg correction tests. The only significant difference (**\*** indicates *p* ≤ 0.05) was observed for the phylogenetic diversity (Faith’s PD) between seawater and whale’s respiratory vapour samples.

| Comparison | Metric | Chi Square | DF | P value KW | P value PW (BH) |
| --- | --- | --- | --- | --- | --- |
| Air vs Whales | Richness | 1.35 | 1 | 0.24 | 0.44 |
| Air vs Whales | Shannon | 2.4 | 1 | 0.12 | 0.22 |
| Air vs Whales | Simpson | 2.4 | 1 | 0.12 | 0.22 |
| Air vs Whales | Faith | 0.60 | 1 | 0.43 | 0.66 |
| Seawater vs Whales | Richness | 3.34 | 1 | 0.06 | 0.08 |
| Seawater vs Whales | Shannon | 0.61 | 1 | 0.43 | 0.53 |
| Seawater vs Whales | Simpson | 0.27 | 1 | 0.60 | 0.71 |
| Seawater vs Whales | Faith | 4.36 | 1 | **0.04\*** | **0.04\*** |
| Species vs Species | Richness | 3.12 | 2 | 0.20 | All $\geq 0.5$ (Bps v Mns v Bms) |
| Species vs Species | Shannon | 2.45 | 2 | 0.29 | All $\geq 0.6$ (Bps v Mns v Bms) |
| Species vs Species | Simpson | 3.16 | 2 | 0.20 | All $\geq 0.4$ (Bps v Mns v Bms) |
| Species vs Species | Faith | 2.45 | 2 | 0.29 | All $\geq 0.6$ (Bps v Mns v Bms) |

### 4.2 Beta diversity: whale blow vs seawater

Beta-diversity analyses based on Bray-Curtis dissimilarities of relative abundances revealed a significant effect of environment (whale blow vs. seawater) on community composition (PERMANOVA: *R*^2^ = 0.140, *F* = 1.31, *p* = 0.030). Consistent results were obtained using Jaccard dissimilarities based on presence-absence data, which also indicated significant differences between environments (PERMANOVA: *R*^2^ = 0.118, *F* = 1.07, *p* = 0.019).

Phylogenetic differences between whale blow and seawater microbial communities were further examined using weighted and unweighted UniFrac distances. Both UniFrac metrics detected significant differences between environments in PERMANOVA analyses, and tests for homogeneity of multivariate dispersion revealed significantly greater within-group variability in seawater samples compared with whale blow samples (PERMDISP: unweighted UniFrac, *F* = 52.9, *p* = 0.001; weighted UniFrac, *F* = 11.2, *p* = 0.018) (Figure 3).

**Figure 3.**
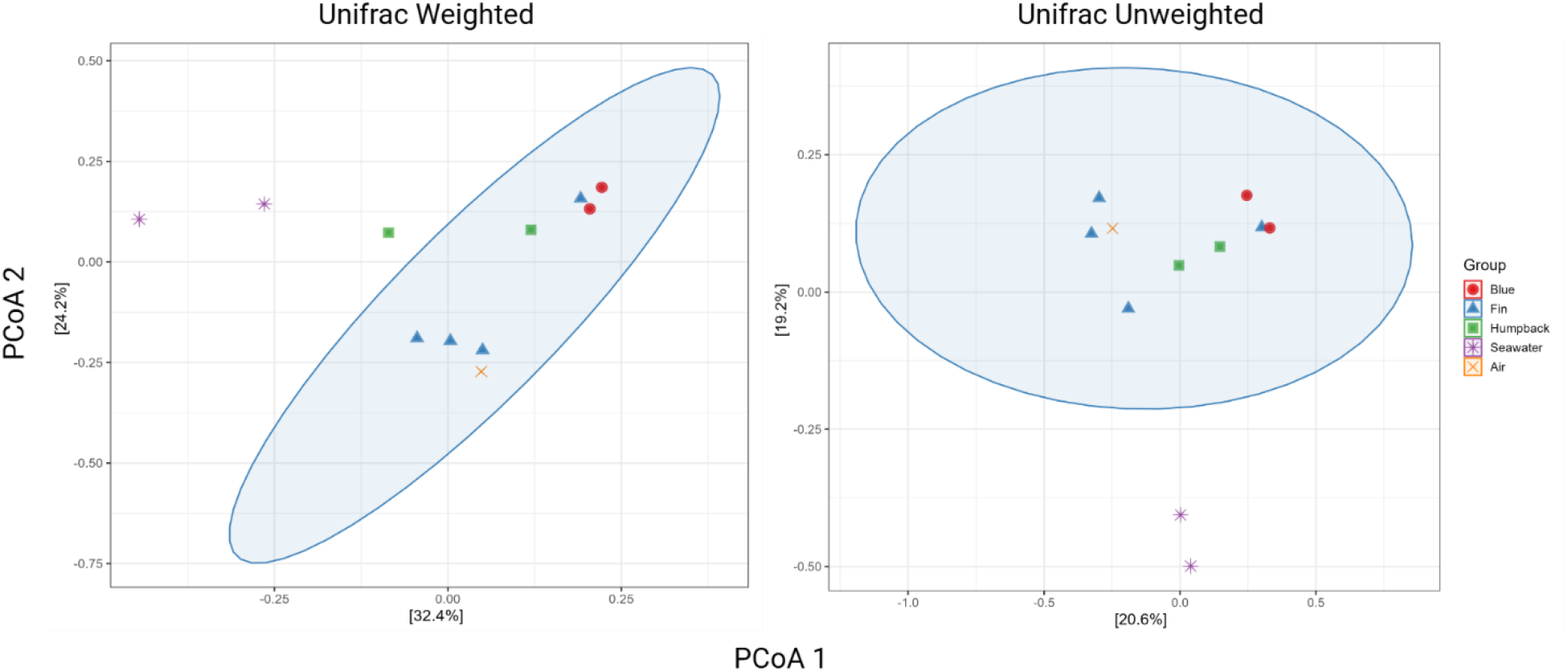
Unifrac scores showing significant differences between whales and seawater for weighted (*F*= 11.2, *p* = 0.018) and unweighted (*F*=52.9, *p* = 0.001). Confidence Ellipses (95%) show the expected dispersion in multivariate spaces.

### 4.3 Phylogenetic beta-diversity among whale species

Within whale blow samples, phylogenetic beta-diversity did not differ significantly among fin, humpback, and blue whales when assessed using either unweighted or weighted UniFrac distances (PERMANOVA: unweighted UniFrac, *R*^2^ = 0.266, *F* = 0.91, *p* = 0.626; weighted UniFrac, *R*^2^ = 0.226, *F* = 0.73, *p* = 0.815). Tests for homogeneity of multivariate dispersion revealed modest differences in within-group variability among species for unweighted UniFrac (PERMDISP, *p* = 0.025), whereas no differences in dispersion were detected for weighted UniFrac (PERMDISP, *p* = 0.461).

### 4.4 Bacterial community characteristics in whales

A total of 601 unique ASVs were assigned to 24 phyla, 37 classes, 84 orders, 124 families and 157 genera in the whale respiratory blow samples. The core microbiota shared by all whales and all samples comprised six phyla (Pseudomonadota, Bacteroidota, Bacillota, Verrucomicrobiota, Cyanobacteriota and Patescibacteria), five classes (Gammaproteobacteria, Bacteroidia, Alphaproteobacteria, Verrucomicrobiia and Clostridia), three orders (Burkholderiales, Pseudomonadales and Enterobacterales) and one family (Comamonadaceae).

However, when considering the 50 most abundant ASVs (relative abundance ≥ 0.05% in at least one whale), the dataset comprised 10 phyla associated with 25 genera (Figure 4). Pseudomonadota was the most abundant phylum across all individuals, particularly in fin whales, followed by Verrucomicrobiota. At the genus level, *Psychrobacter* was the most abundant and prevalent taxon, being absent only from samples Bp053-02 and Bp053-03, whereas *Burkholderia* dominated sample Bp053-01, indicating marked intra-individual variation in microbial communities within the same fin whale (Bp053).

**Figure 4.**
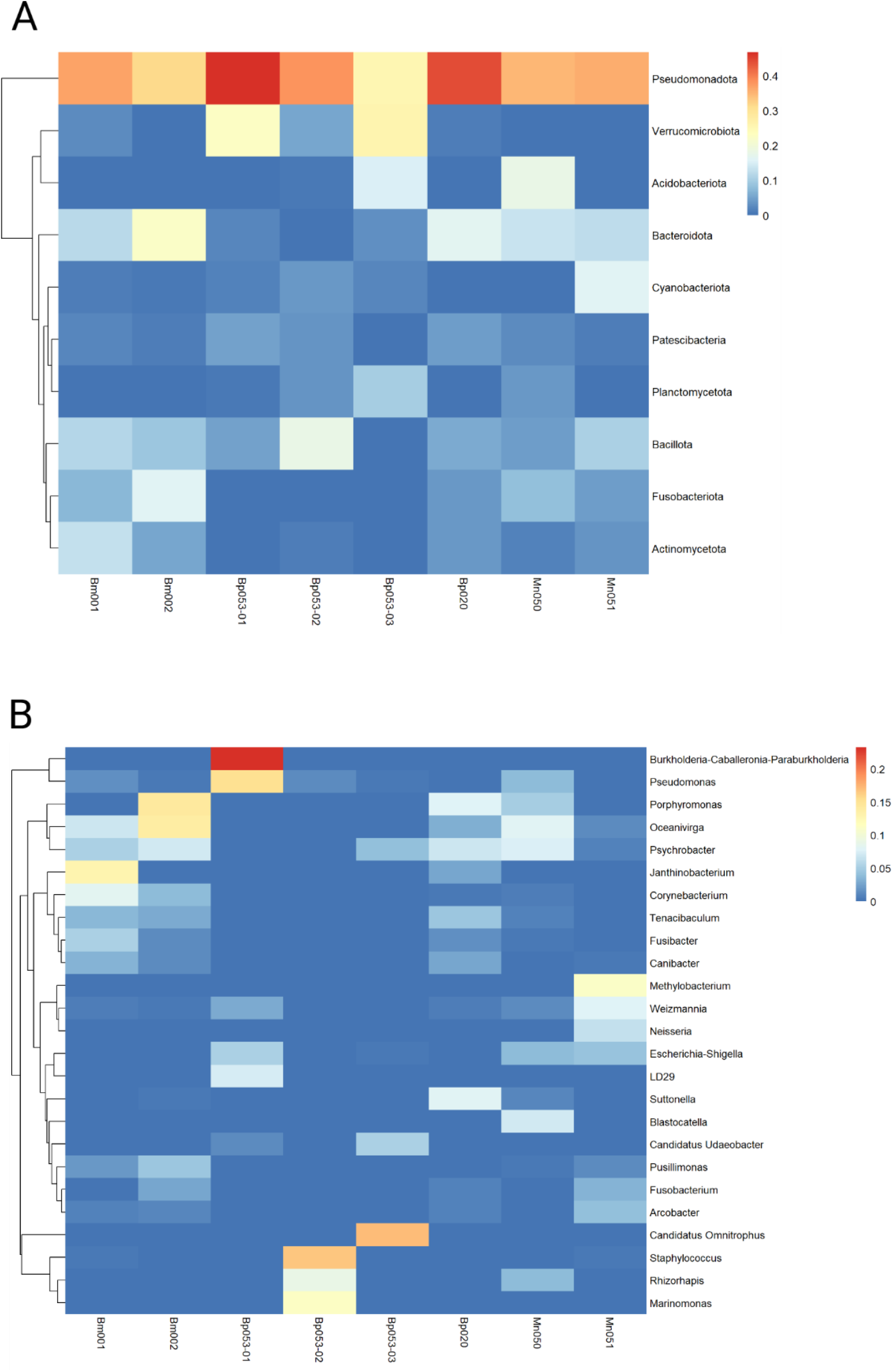
Relative abundance of the most abundant assigned ASVs (top 50), at the phylum (A) and Genus (B) levels for each whale, including the three respiratory samples of whale Bp053 which showed different microbial profiles in each one.

In total, 16 genera (out of 157) previously identified as known or opportunistic pathogens in cetaceans were detected (Table 3). The ratio of commensal to potential pathogenic taxa in each whale was negatively correlated with alpha-diversity metrics (Shannon-Weiner: ρ = −0.88, *p* = 0.01; inverse Simpson: ρ = −0.71, *p* = 0.003; Faith’s PD: ρ = −0.94, *p* = 0.004), indicating that higher relative abundances of potential pathogens were associated with lower alpha diversity (Figure 5). Significant positive correlations were also observed between alpha-diversity metrics (Shannon, inverse Simpson and Faith’s PD) and skin condition scores (ρ = 0.84, *p* = 0.03), suggesting that higher alpha diversity was associated with better skin health, whereas no significant relationships were detected with body condition, injury condition or parasite load.

**Table 3.**
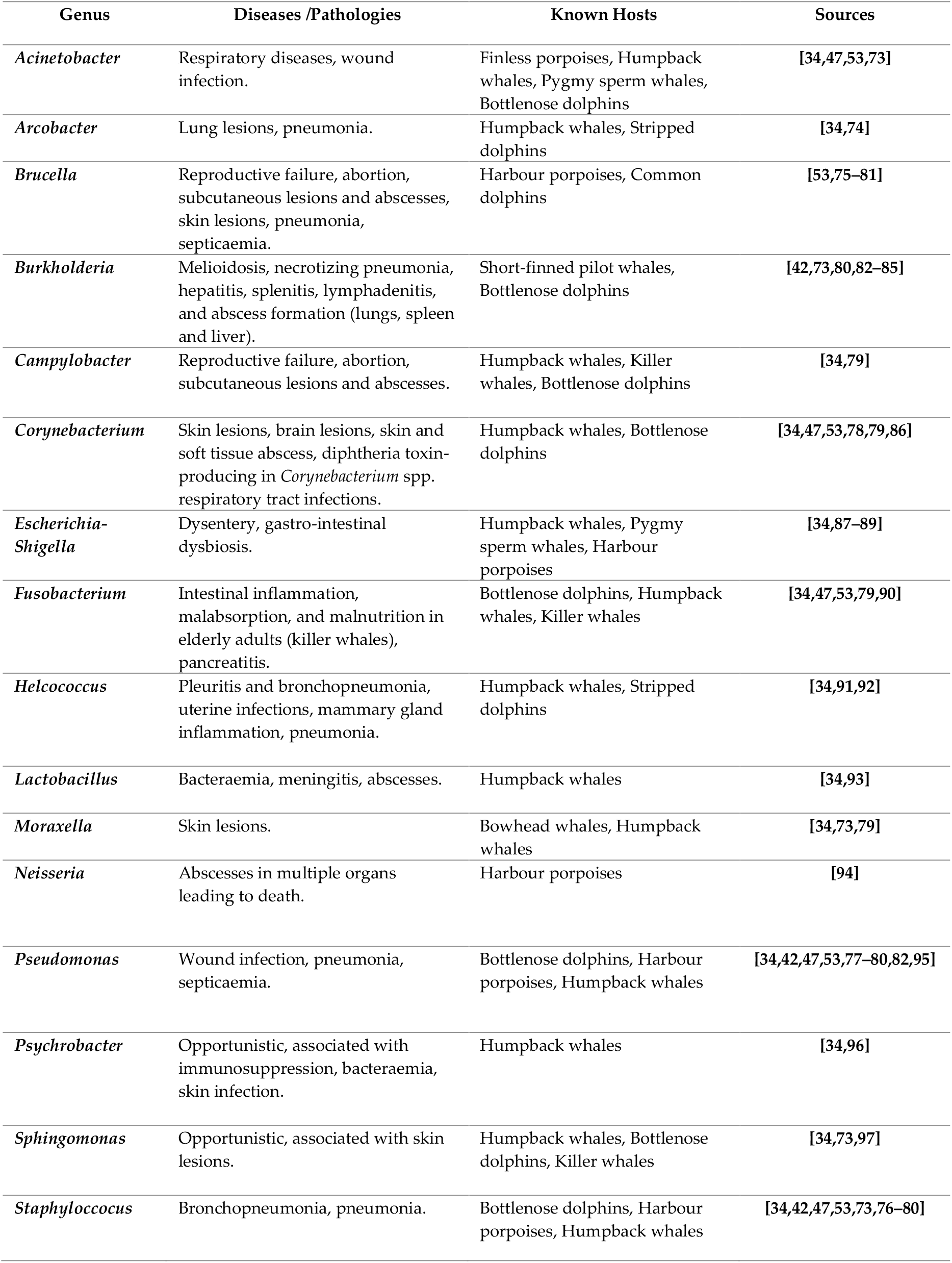
Genera including species of confirmed and opportunistic pathogens found in cetaceans and associated diseases (cetaceans/ mammals). Some sources only mentioned the presence of these genera without any clear evidence of their role in the health status of the host.

**Figure 5.**
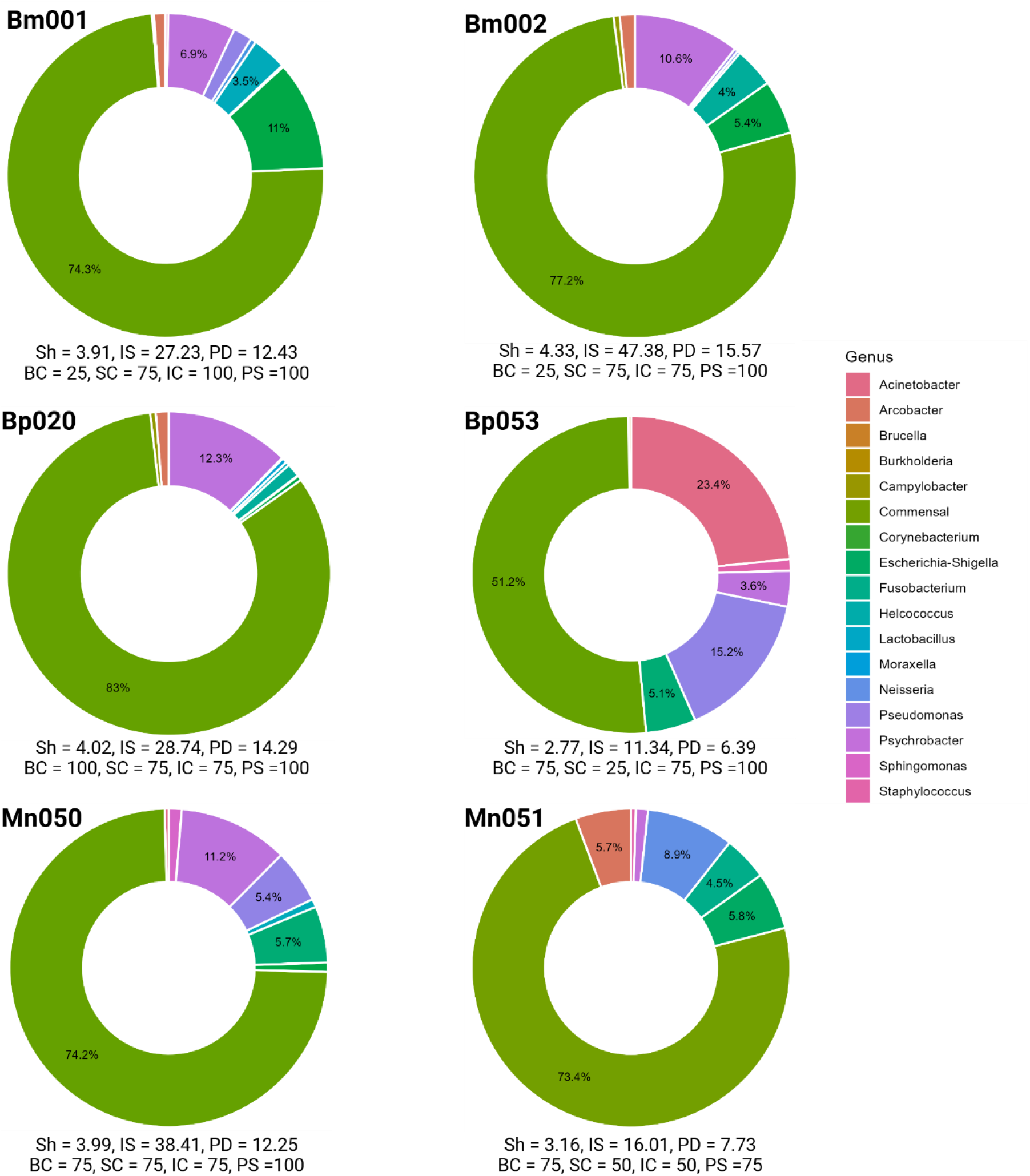
Relative abundance (at the genus level) of commensal (highest % in the green portions) and potential pathogens, for each whale (blue whales: Bm001 and Bm002; fin whales: Bp053 and Bp020, and humpbacks: Mn050 and Mn051). Under the donut pie charts, alpha diversity metrics (Sh = Shannon; IS = Inverse Simpson and PD = Faith’s phylogenetic diversity) and physical health metrics (BC = body condition; SC = skin condition; IC = injury condition and PS = parasite loads) are presented for each individual where the lowest scores relate to the lowest health/welfare states (possible range 0-100).

## 5. Discussion

### 5.1 Diversity metrics

Our analysis of the respiratory microbiota and physical health parameters of six rorqual whales in an anthropised environment (GSL) highlights the potential of these measures as health biomarkers. Although taxonomic alpha-diversity metrics showed limited differences between whale species and seawater samples, Faith’s phylogenetic diversity revealed a significant reduction in evolutionary breadth in intra-whale-associated microbial communities. This phylogenetic signal was mirrored in beta-diversity analyses, with whale and seawater samples showing clear separation in UniFrac space (Figure 3). Taken together, these results suggest that the whale respiratory microbiota represent a phylogenetically structured subset of the surrounding seawater community, consistent with host-associated microbial filtering rather than passive environmental sampling.

### 5.2 Core bacterial taxonomy

Of the 10 most abundant phyla identified in our whales’ respiratory vapour, and not in surface seawater samples, nor in the air sample, eight were previously identified as core bacterial taxa in the exhaled breath of cetaceans: Pseudomonadota (synonym: Proteobacteria) [34,40,43–45,50], Bacteroidota [34,40,43–45,47,50], Fusobacteriota [34,40,44,50], Cyanobacteriota (synonym: Oxyphotobacteria)[40], Patescibacteria [40,44], Bacillota (synonym: Firmicutes)[34,40,43–45,50], Fusobacteriota [34,40,44,50], and Actinomycetota (synonym: Actinobacteria) [34,40,43–45,51]. Whereas Verrucomicrobiota was associated with gut and skin bacteriome of other cetacean species [98,99], Acidobacteriota was not, to our knowledge, reported in any cetaceans or other marine mammal species. At the genus level, our results also aligned with previous studies characterizing the exhaled breath bacteriome of cetaceans, notably the genera *Oceanivirga, Tenacibaculum*, and *Porphyromonas* which have been previously found in blue [50], humpback [34], and North Atlantic right whales (*Eubalaena glacialis*) [100]. *Methylobacterium* and *Suttonella*, have also previously been found in multiple species of free-ranging and captive cetacean respiratory tracts [40,44,50,101–104]. Other genera previously observed in cetaceans included *Janthinobacterium*, a genus associated with cold environments and found on the skin microbiome of humpback and beluga whales (*Delphinapterus leucas*) [105,106], and *Fusibacter*, found in the oral cavity of a stranded pygmy sperm whale (*Kogia breviceps*) [104]. Interestingly, the genus *Canibacter*, highly abundant in the two blue whales and humpback Mn053 presented here, is associated with the normal oral flora of domestic dogs (*Canis familiaris*) but was recently found in infected bite wounds in humans [107]. Since *Canibacter* is only considered an emerging pathogen in humans, we did not include this genus in our pathogen category. However, Mn053 also harbored *Neisseria* in his respiratory tract, a genus commonly found in the oral flora of domestic dogs and grey seals (*Halichoerus grypus*), and previously implicated in fatal bite wound infections in harbor porpoises (*Phocoena phocoena*). Together, these findings highlight the need for further investigation of *Neisseria* and *Canibacter* in marine mammals, particularly in the oral cavity of grey seals and during necropsies of harbor porpoises. Finally, the genera *LD-29* and *Marinomonas* were found to be associated with bacterioplankton and microalgal microbiome [108,109], whilst *Blastocatella, Candidatus Udaeobacter, Pusillimonas, and Candidatus Omnitrophus* were associated with soil and pharmaceutical wastewater [110,111], grassland soil and antibiotic resistance [112–114], landfill leachate [115] and aquatic ecosystems undergoing groundwater salinization disruption [116,117] respectively. Importantly, these last genera, which interestingly were not found in the surrounding seawater samples, reflect the well documented eutrophication of the GSL. The fact that some of these bacterial taxa have been previously associated with antibiotic resistance is concerning, especially for the recovery of blue whales, which are endangered in Eastern Canada with an estimated number of approximately 250-400 individuals [68]. In such small and vulnerable populations, even subtle additional health stressors can have important consequences at both individual and population levels [118]. Antibiotic-resistant bacteria carry resistance determinants in their genes, allowing them to persist in the environment and facilitating the spread of these resistant traits to other bacteria through horizontal genetic exchange [119,120]. Beyond limiting therapeutic options, the dissemination of resistance can restructure microbial communities and favour the emergence of opportunistic pathogens with greater pathogenic potential. Importantly, resistance mechanisms are not just defensive, they often entail changes in bacterial physiology, regulation, and metabolism that can enhance virulence [119]. This resistance–virulence interplay is characterised by an increased capacity for colonisation, immune evasion, and persistence within the host, ultimately leading to more severe infections. In the context of endangered blue whales already exposed to multiple anthropogenic stressors, the presence of such microbial risks raises additional concerns for individual welfare and long-term population resilience.

### 5.3 Taxonomic differences in relation to respiratory events

Fin whale ID Bp053 was sampled three times during one ventilation sequence (duration of 2 min 53sec at the surface) which comprised 12 respiratory events (spouts). Although the bacterial profiles of the three samples differed taxonomically, no significant differences were observed in the diversity metrics (Table1 and Figure 4). Interestingly, *LD-29* and *Marinomonas* genera were found only in the first sample (Bp053-01) at the beginning of the ventilation sequence. The genus *LD-29* as we described earlier, is found in bacterioplankton and *Marinomonas* in microalgal specimen but was not found in our seawater surface layer samples. This result could be associated with the underlying feeding mechanisms and anatomy of rorqual whales: during feeding, an elevation of the oral plug shuts the oropharyngeal channel for allowing the pharynx to be dedicated to prey ingestion and then conversely shuts down to open the airways. Since rorqual whales are in the GSL to feed, *LD-29* and *Marinomonas* might have been transferred into the upper respiratory tract when the oral plug shifted after feeding [32]. The differences in genera found in the second and third respiratory samples might be associated with the bacterial ecological niches either in the upper or lower respiratory tract, as it is the case in many animals [121,122]. Moreover, the tidal volume of cetacean ventilation cycles is greater than terrestrial animals, which could yield a greater diversity of expelled bacteria from both upper and lower respiratory tracts [123]. Our results suggest that though microbial diversity is generally consistent from breath to breath in cetaceans, there may be alterations in the probability of sampling certain taxa from the beginning to the end of a surfacing sequence, and therefore we recommend further research on this unique observation.

### 5.4 Airway pathobionts and physical health measures

To our knowledge, the current study is the first to examine the potential relationships between whale health parameters (body condition, skin health, injury condition and ectoparasite load scores) [66], alpha-diversity metrics, and relative abundance of opportunistic and true pathogens in exhaled breath. Whales with lower alpha diversity metrics had the highest pathogen abundance in their respiratory microbiota. For instance, fin whale Bp053 had the lowest Shannon-Weiner index of all individuals, with a score of 2.77, and 48.8% of her microbiota was associated with pathobionts (including the genus *Brucella*, a well documented pathogen in cetaceans) indicating reduced diversity and evenness of the microbial community and microbial imbalances (dysbiosis). In comparison, blue whale Bm002, which had the highest Shannon-Weiner score of 4.33 and only 22.8% abundance of pathobionts in her respiratory microbiota, points towards a richer and more even microbial community. In terrestrial mammals, research on airway pathobiomes and their relationships with alpha diversity metrics is a relatively recent field [1]. Some studies, notably in porcine, bovine, canine and feline species, found significant differences between healthy and diseased animals, with reduced alpha-diversity observed in unhealthy animals [22,29], whereas others did not find statistical significant differences between the two groups [124,125]. Multiple factors can influence a host’s microbial community, including age, environmental conditions, and food sources, and these variables should be systematically included in research protocols. In our study, we also included physical health measures (body condition, skin condition, known past and present injuries, and ectoparasite load) and found that higher alpha-diversity, as in Bp020, Bm001, and Bm002, was correlated with better skin health. Future studies should include larger sample sizes of individual whales and more variables, such as seawater physicochemical properties, prey availability, and animal life status (age), although these can be difficult to determine in free-ranging cetaceans.

## 6. Conclusion

Our study has established a first baseline for the respiratory microbiota of rorqual whales in the Gulf of St-Lawrence and highlights the potential of alpha-diversity metrics, abundance of pathogens, and whale physical indicators, notably skin condition, as biomarkers of health. Nevertheless, we point out the limitations of this study regarding the small sample size and opportunistic nature of sampling. Given the conservation status of blue and fin whales in Eastern Canada, and the increasing anthropogenic pressures on this ecosystem, such integrative, non-invasive monitoring tools can become valuable components of long-term health and welfare surveillance strategies in the conservation of these species.

## Acknowledgments

We would like to thank Robert Michaud (GREMM) for sharing available information about life stages of the 6 whales; Jacques Gélineau, Larry Mercier, Caroline Demontigny, Véronique Demontigny, Julie Noel and Yves Jean for sharing whale sightings. Mario Guay for laboratory work. We would also like to thank the Cégep de Sept-Îles for its continuous support.

## Ethical Statement

The current research follows the Canadian Council on Animal Care and received ethical approval from the Animal Care Committee of the University Laval (2020-2025-433, VRR-24-433) and the Division of Fisheries and Oceans Canada (research permit QUE-LEP-014B-2020-2025).

### Authors’ Contributions

Conception of the study (AB, MD, JAD); Data acquisition (AB, JB, AM, RP, MC); Data analysis and interpretation (AB, CV, JAD, AM, KH); Drafting and revision (AB, CV, JAD, AM, JB, MD, MC and KH).

